# Minimum Effective Dose of Clemastine in a Mouse Model of Preterm White Matter Injury

**DOI:** 10.1101/2024.02.08.578953

**Authors:** Elizabeth Odell, Nora Jabassini, Björn Schniedewind, Sarah E. Pease-Raissi, Adam Frymoyer, Uwe Christians, Ari J. Green, Jonah R. Chan, Bridget E.L. Ostrem

## Abstract

**Background:** Preterm white matter injury (PWMI) is the most common cause of brain injury in premature neonates. PWMI involves a differentiation arrest of oligodendrocytes, the myelinating cells of the central nervous system. Clemastine was previously shown to induce oligodendrocyte differentiation and myelination in mouse models of PWMI at a dose of 10 mg/kg/day. The minimum effective dose (MED) of clemastine is unknown. Identification if the MED is essential for maximizing safety and efficacy in neonatal clinical trials. We hypothesized that the MED in neonatal mice is lower than 10 mg/kg/day.

**Methods:** Mouse pups were exposed to normoxia or hypoxia (10% FiO_2_) from postnatal day 3 (P3) through P10. Vehicle or clemastine fumarate at one of four doses (0.5, 2, 7.5 or 10 mg/kg/day) was given orally to hypoxia-exposed pups. At P14, myelination was assessed by immunohistochemistry and electron microscopy to determine the MED. Clemastine pharmacokinetics were evaluated at steady-state on day 8 of treatment.

**Results:** Clemastine rescued hypoxia-induced hypomyelination with a MED of 7.5 mg/kg/day. Pharmacokinetic analysis of the MED revealed C_max_ 44.0 ng/mL, t_1/2_ 4.6 hours, and AUC_24_ 280.1 ng*hr/mL.

**Conclusion:** Based on these results, myelination-promoting exposures should be achievable with oral doses of clemastine in neonates with PWMI.

**Key Points:**

- Preterm white matter injury (PWMI) is the most common cause of brain injury and cerebral palsy in premature neonates.
- Clemastine, an FDA-approved antihistamine, was recently identified to strongly promote myelination in a mouse model of PWMI and is a possible treatment.
- The minimum effective dose in neonatal rodents is unknown and is critical for guiding dose selection and balancing efficacy with toxicity in future clinical trials.
- We identified the minimum effective dose of clemastine and the associated pharmacokinetics in a murine chronic hypoxia model of PWMI, paving the way for a future clinical trial in human neonates.

## INTRODUCTION

Preterm white matter injury (PWMI) affects approximately 20,000 neonates in the United States annually and can result in lifelong motor and cognitive disability^1–3^. There are no specific treatments available; symptom-directed supportive care is the mainstay of management^4^. Oligodendrocytes (OLs) and their precursors (oligodendrocyte precursor cells, OPCs) comprise the major cell types implicated in PWMI, which involves an arrest of differentiation of OPCs and a reduction in mature OLs and myelin formation^5–11^. The OL lineage is therefore an ideal target for therapeutics aimed at promoting recovery in the setting of PWMI.

A previous high-throughput screen using a novel micropillar array-based assay identified several FDA-approved compounds, including the antihistamine clemastine, that promote differentiation of OPCs into mature, myelinating OLs^12^. Clemastine at doses of 10 mg/kg/day was further demonstrated to enhance myelination and functional recovery in several animal models of white matter injury, including PWMI^13–19^. However, the minimum effective dose (MED) that promotes neonatal brain repair in mice is unknown. Identification of the MED is essential to guide therapeutic development and balance efficacy with toxicity and unnecessary side effects, especially in a neonatal population. A pharmacokinetic (PK) understanding of the MED in mice can further aid in determining the optimal dosing scheme and target exposure for Phase I studies in humans.

Here, we use a chronic hypoxia model in neonatal mice that recapitulates major features of PWMI, including diffuse hypomyelination, OL maturation delay and persistent motor and cognitive deficits^18–20^. From postnatal day 3 (P3) through P10, mouse pups are subjected to 7 days of hypoxia, mimicking a known risk factor for PWMI^21^. When considering equivalent stages of brain development and myelination, P3 through P10 in mice corresponds to gestational weeks 23 through 40 in human neonates: the highest risk period for PWMI^22–24^. Using the murine chronic hypoxia model, we determine the MED of clemastine, which we defined as the lowest dose that promoted myelination above vehicle control levels by all measures tested. We characterized the pharmacokinetics of the MED, paving the way for a future clinical trial of clemastine in neonates with PWMI.

## METHODS

### Chronic hypoxia and clemastine treatment

All animal studies were approved by the University of California, San Francisco, Institutional Animal Care and Use Committee. Male and female C57BL/J mice were used in equal proportions for all experiments except where indicated otherwise. Mice were housed in temperature- and humidity-controlled environments on a 12h/12h light/dark cycle with free access to standard chow and water. For chronic hypoxia experiments, mouse pups with their lactating mothers were subjected to chronic sublethal hypoxia (10% fraction of inspired oxygen [FiO_2_]) from P3 through P10. Mouse pups were treated daily from P3 through P10 with vehicle (saline) or clemastine fumarate (Selleckchem) by oral gavage. Doses of clemastine (weighed and dosed based on the clemastine fumarate salt) were: 0.5, 2, 7.5, or 10 mg per kg of body weight. On P10, mice were returned to normoxic (21% FiO_2_) conditions. At P14, OL differentiation and myelination were compared in hypoxic mice treated with vehicle or clemastine and in mice exposed to normoxia from P0 through P14.

### Immunohistochemistry

P14 mice were euthanized and perfused transcardially with ice-cold Phosphate Buffered Saline (PBS) followed by ice-cold 4% paraformaldehyde (Electron Microscopy Sciences) diluted in water. Brains were removed and stored overnight in 4% paraformaldehyde at 4°C. Tissues were then placed in 30% sucrose in PBS at 4°C for 1-2 days followed by freezing in O.C.T. compound (Tissue-Tek) and generation of 30 um coronal brain sections on a microtome (HM 450 Sliding Microtome, Epredia™, Richard-Allan Scientific). Sections were blocked and permeabilized for 2 hours at room temperature in blocking solution (PBS with 0.1% Triton X-100, and 10% donkey or goat serum), and subsequently incubated overnight at 4°C in blocking solution with primary antibody added. After washing, secondary antibody incubation was performed for 2h at room temperature in 10% donkey or 10% goat serum in PBS with secondary antibody added. Primary antibodies used were: rat anti-MBP (1:500, MCA409S, Serotec), mouse anti-Olig2 (1:200, EMD Millipore, MABN50), rabbit anti-Cleaved Caspase-3 (CC3, 1:300, Cell Signaling 9661S), and rabbit anti-Sox10 (1:500, EMD Millipore AB 5727). Secondary antibodies were: donkey or goat AlexaFluor 488 (1:1,000), 594 (1:1000), or 647 (1:500)-conjugated IgG. MBP fluorescence signal intensities were measured in a fixed area of striatum using Imaris software, normalized within each coronal brain slice to the area of lowest fluorescence. Cell counts for Sox10-, CC1-, and CC3-positive cells were performed using Imaris software, Count Spots function, manually adjusted to remove incorrect spots (for example double counting of single cells or counting of fluorescent debris). All fluorescence intensity quantifications and cell counting were performed by an investigator blinded to the experimental condition.

### Electron microscopy

P14 mice were euthanized and perfused transcardially with ice-cold 0.1M sodium cacodylate buffer (0.1M sodium cacodylate trihydrate [Electron Microscopy Sciences, 12310], 5 mM calcium chloride dihydrate [Sigma, 223506], pH adjusted to 7.3-7.4), followed by ice-cold EM fixative (1.25% glutaraldehyde [Electron Microscopy Sciences, 16220], 2% paraformaldehyde [Electron Microscopy Sciences, 19210], 0.1M sodium cacodylate buffer). Brains were removed and stored for 8 days in EM fixative at 4°C, and subsequently placed in 30% sucrose in PBS at 4°C for 1-2 days. Samples were mounted in O.C.T. compound (Tissue-Tek) and 500 um coronal sections were generated on a microtome (HM 450 Sliding Microtome, Epredia™, Richard-Allan Scientific). The section at approximately +1.1 mm anterior to bregma (joining of corpus callosum) was placed in PBS and under a dissecting microscope, the middle 1/3 of the corpus callosum was dissected using a razor blade. Samples were stained with osmium tetroxide, dehydrated in ethanol and embedded in TAAB resin. Sections were cut perpendicular to the angle of the fibers of the corpus callosum at 1-mm intervals and stained with toluidine blue for light microscopy analysis. Axons were then examined using electron microscopy, and g-ratios were calculated as the diameter of the axon divided by the diameter of the axon and the surrounding myelin sheath. We could not reliably identify unmyelinated axons in the corpus callosum at P14, even at high resolution (11.5K magnification). Thus, all quantifications were performed on myelinated axons only. Measurements were performed by an investigator blinded to the experimental condition.

### Plasma sample collection and mass spectrometry

Male and female C57BL/J mice were dosed by oral gavage with 7.5 mg/kg/day clemastine daily from P3 to P10. Blood was sampled before (time=0) the 8^th^ dose and at 0.5, 1, 1.5, 2, 3, 4, 6, 9, 12, and 24h after the 8^th^ dose. Terminal blood collections were performed by cardiac puncture after deeply anesthetizing the animal. At least 3 animals were sampled per timepoint and at least one animal per sex was included for every time point. Blood was collected into 1 ml syringes using 25ga needles coated with 0.5M EDTA (ThermoFisher, #15575020), and immediately gently transferred into K2EDTA tubes (Sarstedt #41.1395.105) followed by centrifugation at 6,700 rcf for 5 minutes at 4°C. Plasma was transferred into cryotubes (Nalgene™ Cryogenic Tube, #5012-0020) and stored at −80°C until sample analysis.

Clemastine was quantified in mouse EDTA plasma using high-performance chromatography-tandem mass spectrometry (LC-MS/MS) at iC42 Clinical Research and Development (University of Colorado, Aurora, CO, USA). The assay followed the principles described by Xie *et al*^25^. Clemastine reference material was from Toronto Research Chemicals (North York, ON, Canada) and the internal standard diphenhydramine-D_3_ from Sigma Aldrich (St. Louis, MO, USA). Isotope-labeled clemastine as internal standard was commercially not available at the time of the study. Two hundred (200) µL of a protein precipitation solution (0.2 M ZnSO_4_ 30% water/ 70% methanol v/v) containing the internal standard (1.0 ng/mL diphenhydramine-D_3_) was added to 50 µL of study samples, quality control samples, calibrators and zero samples. Samples were vortexed for 2.5 min, centrifuged at 4°C and 16,000 g for 10 min. The supernatants were transferred into 2 mL glass HPLC injection vials. The samples were then further extracted online and analyzed using a 2D-LC-MS/MS system composed of Agilent 1100 HPLC components (Agilent Technologies, Santa Clara, CA USA) and a Sciex API 5000 MS/MS detector (Sciex, Concord, ON, Canada) connected via a turbo flow electrospray source run in the positive ionization mode (4500V, 550°C source temperature).

Ten (10) µL of the samples were injected onto the extraction column (Zorbax XDB C8, 4.6 · 50 mm, Agilent Technologies). The mobile phase was 80% 0.1% formic acid in HPLC grade water (mobile phase A) and 20% methanol containing 0.1% formic acid (mobile phase B). Samples were cleaned with a solvent flow of 3 mL/min and the temperature for the extraction column was set to room temperature. After 0.7 min, the switching valve was activated and the analytes were eluted in the backflush mode from the extraction column onto a 4.6 · 150 mm analytical column filled with C8 material of, 5 μm particle size (Zorbax XDB C8, Agilent Technologies). The analytes were eluted using a gradient starting with 50% mobile phase B that increased to 98% within 2.3 min and was held for 1.0 min. The system was then re-equilibrated to starting conditions at 50% B for 0.8 min. The flow rate was 1.0 mL/min and the analytical column was kept at 60°C. The MS/MS was run in the multiple reaction mode (MRM) and the following ion transitions were monitored: clemastine *m/z*= 346.2 [M(^37^Cl) +H]^+^ → 217.0 (quantifier), *m/z*= 344.2 [M(^35^Cl) +H]^+^ → 215.0 (qualifier) and diphenhydramine-D_3_ (internal standard) *m/z*= 259.0 [M+H]^+^ → 167.3. Delustering potentials were set to 51V and collision energies were set to 23V for clemastine quantifier and qualifier transitions. For the internal standard diphenhydramine-D_3_ the declustering potential was 56V and the collision energy 19V.

Clemastine concentrations were quantified using the calibration curves that were constructed by plotting nominal concentration *versus* analyte area to internal standard area ratios (response) using a quadratic fit and 1/x weight. All calculations were carried out using the Sciex Analyst Software (version 1.7.3). The quantification range for clemastine was 0.0025 (lower limit of quantification) – 20.0 ng/mL and study sample were diluted 1:50 and 1:250 as necessary for the detector response to fall within the calibration range as necessary. All results reported were from runs that met the following acceptance criteria: >75% of the calibrators had to be within ±15% of the nominal values (except at the lower limit of quantification: ±20%) and >2/3 of the quality controls had to be within ±15% of the nominal values. The imprecision of the results was better than 15%. Significant carry-over and matrix effects were excluded and dilution integrity was established.

### Pharmacokinetic analysis

Non-compartmental PK analysis was conducted using the geometric mean of each timepoint. Steady-state conditions were assumed after 7 days of dosing. The maximum concentration (C_max_), time of the maximum concentration (T_max_), and the concentration at 24 hours post dose (C_24h_) were taken directly from the observed data. Area under the curve during the 24 hour dosing interval (AUC_24_) was calculated using the trapezoidal method. Oral Clearance (CL/F) was then calculated as Dose / AUC_24_. The terminal elimination rate constant (k_e_) was calculated using linear regression of log transformed concentrations over the terminal log-linear decline phase and from this the terminal elimination half-life calculated (t_1/2_ = 0.693/k_e_).

### Statistical analysis

Statistical significance between groups was determined with GraphPad Prism 5 software. Statistical analyses were performed by one-way analysis of variance (ANOVA) followed by Tukey’s post hoc test. A probability of p<0.05 was considered statistically significant.

## RESULTS

### Identification of minimum effective dose of clemastine

To determine the lowest oral dose of clemastine that provides the full myelination-promoting effect of the medication, we used a murine chronic hypoxia model. Mouse pups and lactating dams were subjected to chronic sublethal hypoxia (10% fraction of inspired oxygen [FiO_2_]) from postnatal day 3 (P3) through P10 (Fig. 1a). This treatment leads to widespread OPC maturation delay and hypomyelination, mimicking histopathological findings in human neonates^7,8,18,26,27^. Hypoxia-exposed mice were treated once daily by oral gavage with either vehicle or clemastine from P3 through P10 and returned to normoxic (21% FiO_2_ conditions) on P10 (Fig. 1b). The highest clemastine dose used was 10 mg/kg/day, which has previously been reported to rescue hypoxia-induced hypomyelination in the murine chronic hypoxia model^18^, and to induce (re)myelination in models of multiple sclerosis^13^, psychiatric disorders^14,15^, aging^28^ and neurodegenerative disorders^29,30^. We tested a 20-fold lower dose (0.5 mg/kg/day) and two intermediate doses, 2 and 7.5 mg/kg/day. We quantified mature OLs at P14 by immunohistochemistry in hypoxia-exposed animals treated with vehicle, hypoxia-exposed animals treated with 0.5, 2, 7.5 or 10 mg/kg/day clemastine, and in normoxia-exposed animals (Fig. 1c, d). Hypoxia led to a significant reduction in OL density (CC1+ cells/area) as previously described^18–20^. Clemastine doses of 7.5 and 10, but not 0.5 or 2, mg/kg/day significantly increased OL differentiation as compared to vehicle (p<0.05, ANOVA followed by Tukey’s post hoc test). Thus, a dose of at least 7.5 mg/kg/day was required for the OL differentiation-inducing effects of clemastine in neonatal mice.

**Figure 1.**
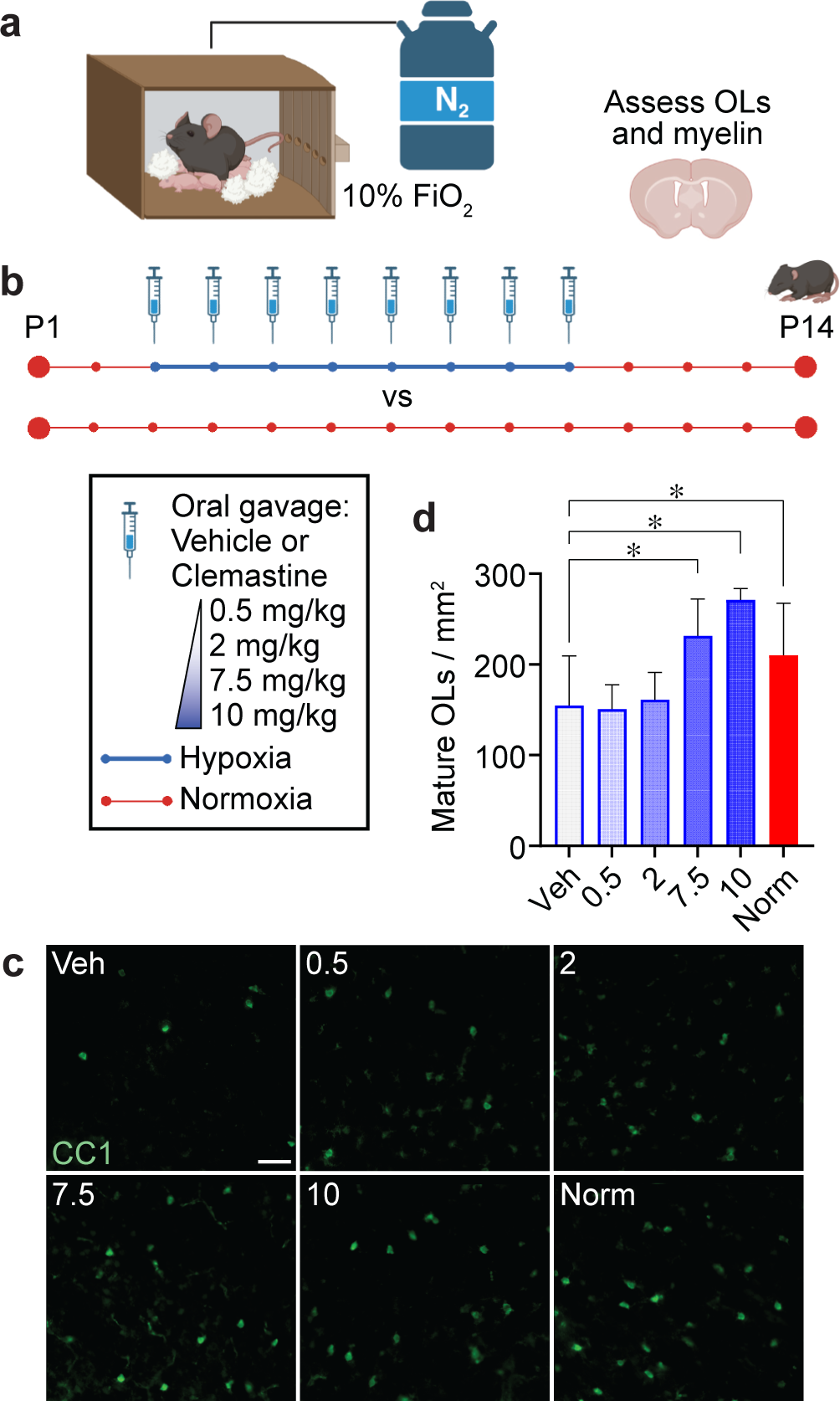
Clemastine rescues hypoxia-induced deficits in oligodendrocyte differentiation. (a) Schematic of hypoxia chamber, in which mice are exposed to 10% FiO_2_ via displacement of oxygen with nitrogen (N_2_) gas. Mice were placed in the chamber in their home cages and given free access to bedding, food and water. (b) Experimental groups and timeline. Mice were exposed to normoxia (21% FiO_2_) throughout the first two weeks of life, or to 10% FiO_2_ from postnatal day 3 (P3) through P10 while undergoing daily treatment with vehicle (saline) or clemastine at 0.5, 2, 7.5, or 10 mg/kg/day. At P14, oligodendrocyte density and myelination were assessed in hypoxic mice treated with vehicle or clemastine and in mice reared in normoxic conditions. (c) Representative images of CC1 staining in the cortex of P14 mice from each treatment condition. Scale bars, 50 um. (d) Quantification of oligodendrocyte density. OLs, oligodendrocytes (CC1+ cells). Hypoxia-exposed animals (blue outline) were: Veh (vehicle-treated), 0.5 (clemastine 0.5 mg/kg/day), 2 (clemastine 2 mg/kg/day), 7.5 (clemastine 7.5 mg/kg/day), 10 (clemastine 10 mg/kg/day). Norm, normoxia-exposed animals (red bar). Error bars represent standard deviation. N= at least 4 mice/condition from at least 3 independent experiments per condition, *p<0.05, ANOVA followed by Tukey’s *post hoc* test. Diagrams were created with BioRender.com.

We assessed myelin via immunohistochemistry in vehicle- and clemastine-treated hypoxia groups and in mice reared in normoxic conditions (Fig. 2a-c). Clemastine at doses of 2 mg/kg/day and above significantly increased myelin basic protein (MBP) intensity at P14 as compared to vehicle levels (p<0.05, ANOVA followed by Tukey’s post hoc test). Given reported sex differences in the incidence and severity of PWMI in humans^31^ and rodents^32^, we assessed for sex differences in OL differentiation and myelination in response to clemastine (Supplementary Fig. 1). We found no significant differences between males and females in any dose group. Accordingly, we did not split mice by sex in our further analyses. In summary, hypoxia-exposed mice treated at 7.5 and 10 mg/kg/day, but not 0.5 or 2 mg/kg/day, had significantly improved myelination and OL differentiation as compared to vehicle-treated hypoxic animals. Based on these results, we established 7.5 mg/kg/day as the MED of clemastine.

**Figure 2.**
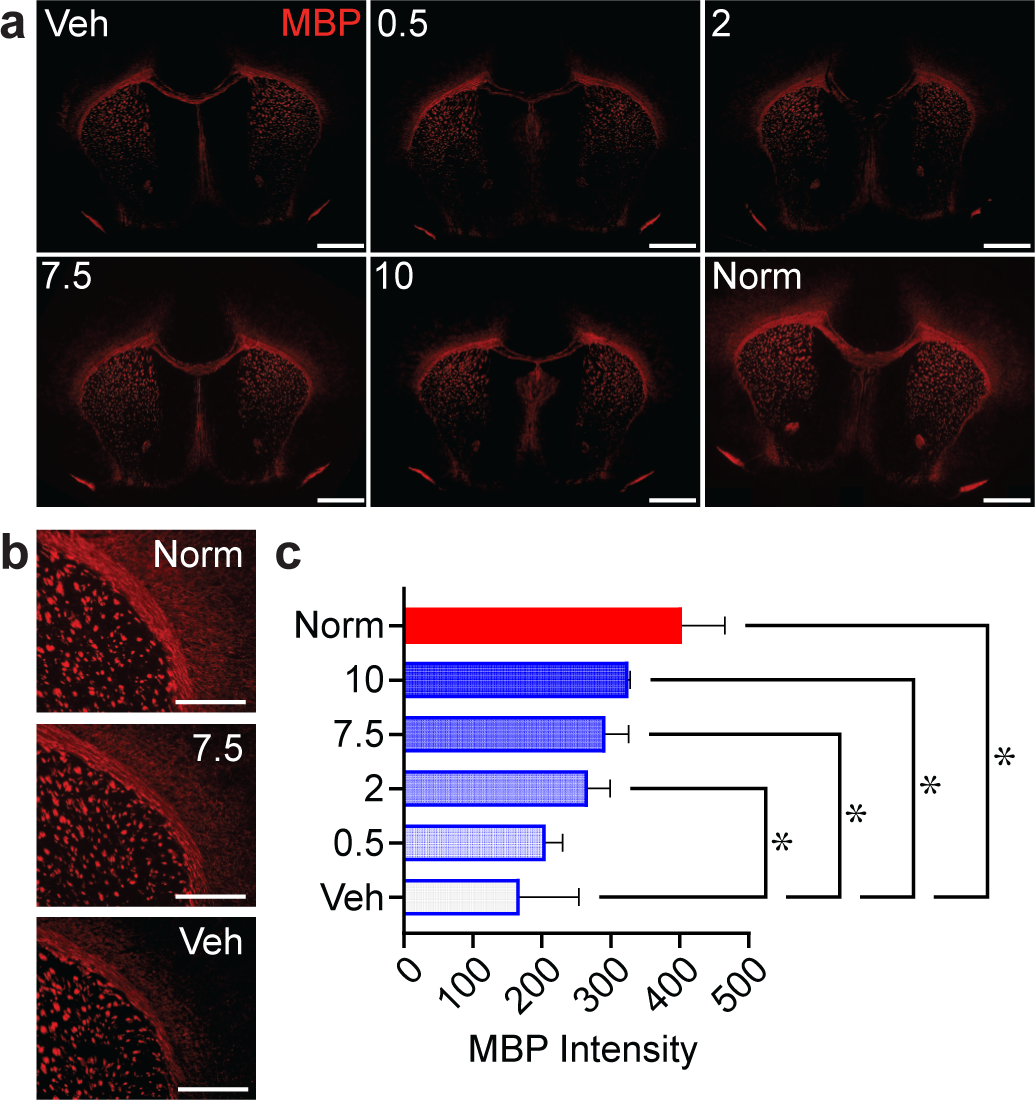
Clemastine improves hypoxia-induced myelination deficits. (a) Representative images of coronal sections stained for myelin basic protein (MBP) at P14. Scale bars, 500 um. (b) Representative images of the corpus callosum and striatum and (c) quantification of staining for MBP in the striatum. N= at least 4 mice/condition from at least 3 independent experiments per condition. *p<0.05, ANOVA followed by Tukey’s *post hoc* test. For all graphs and images, hypoxia-exposed animals were: Veh (vehicle-treated), 0.5 (clemastine 0.5 mg/kg/day), 2 (clemastine 2 mg/kg/day), 7.5 (clemastine 7.5 mg/kg/day), 10 (clemastine 10 mg/kg/day). Norm, normoxia-exposed animals. Error bars represent standard deviation.

We hypothesized that chronic hypoxia leads to an OL differentiation arrest, but not cell death, and that this maturation arrest is rescued by clemastine. We assessed total OL lineage (SOX10+) cells at P14 by immunohistochemistry in hypoxia-exposed animals treated with vehicle or clemastine at 0.5, 2, 7.5 or 10 mg/kg/day and in normoxia-exposed animals (Fig. 3a). Consistent with our hypothesis, total SOX10+ cells were not significantly different across all conditions (Fig. 3b). We calculated the proportion of immature (CC1-) cells (OPCs) out of all OL lineage (SOX10+) cells in each condition (Fig. 3c). The proportion of OPCs was increased in hypoxia-exposed animals treated with vehicle, 0.5 or 2 mg/kg/day clemastine as compared to animals reared in normoxia (p<0.05, ANOVA followed by Tukey’s post hoc test). Thus, chronic hypoxia caused OL lineage cells to shift towards a less mature state, which was rescued by clemastine at doses of 7.5 mg/kg/day and above. We assessed apoptosis in oligodendrocyte lineage cells at P14 by staining for cleaved caspase-3 (Fig. 4a). No significant differences were observed in hypoxia-exposed animals treated with vehicle and clemastine at the MED and in normoxia-exposed animals (Fig. 4b, c). Therefore, hypoxia- and clemastine-induced changes in OL and OPC density and myelination are unlikely to be related to changes in cell death.

**Figure 3.**
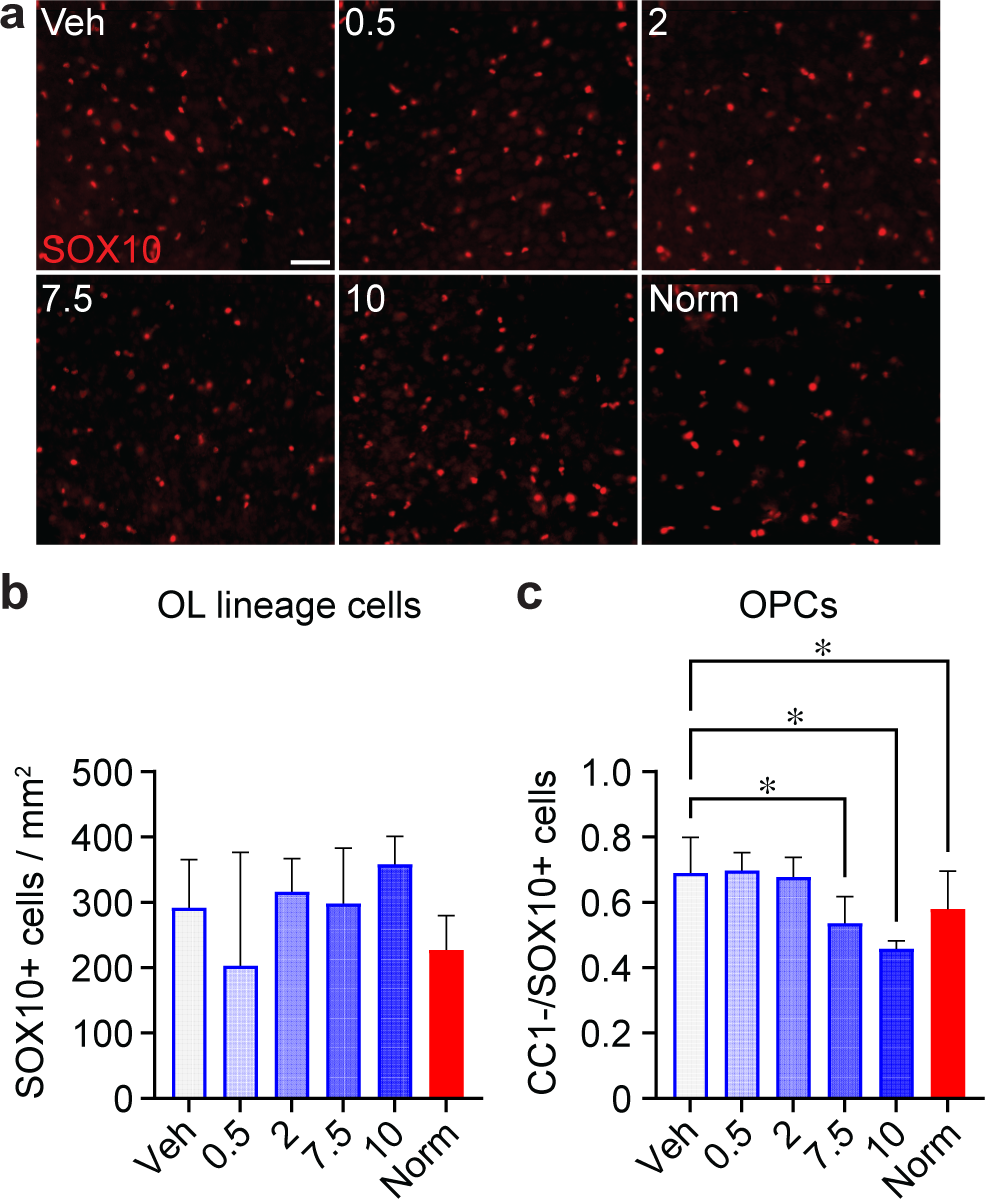
Analysis of the oligodendrocyte lineage in clemastine-treated mice. (a) Representative images and (b) quantification of staining for the pan-oligodendrocyte lineage marker SOX10 in the cortex at P14. Scale bars, 50 um. There were no significant differences between groups. N= at least 4 mice/condition from at least 3 independent experiments per condition. (c) Quantification of the proportion of OPCs (CC1-/SOX10+ cells) out of all OL lineage cells. For all graphs, Hypoxia-exposed animals (blue outline) were: Veh (vehicle-treated), 0.5 (clemastine 0.5 mg/kg/day), 2 (clemastine 2 mg/kg/day), 7.5 (clemastine 7.5 mg/kg/day), 10 (clemastine 10 mg/kg/day). Norm, normoxia-exposed animals (red bars). *p<0.05, ANOVA followed by Tukey’s *post hoc* test. Error bars represent standard deviation.

**Figure 4.**
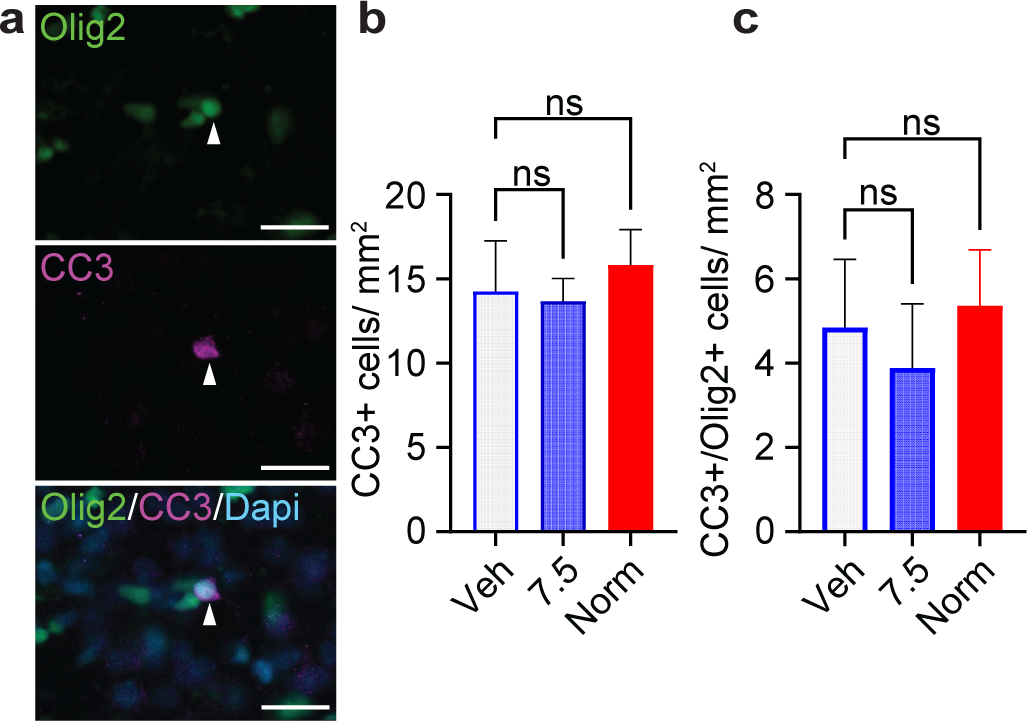
Hypoxia and clemastine treatment do not induce apoptosis in oligodendrocyte lineage cells. (a) Example image of cleaved caspase-3 (CC3)-positive oligodendrocyte lineage (OLIG2+) cell in the periventricular region of a P14 hypoxia-exposed mouse treated with 7.5 mg/kg/day clemastine. Scale bars, 50 um. Apoptotic cells were quantified in coronal brain sections as total CC3+ cells per area (b) and CC3+ oligodendrocyte lineage (CC3+/OLIG2+) cells per area (c) in Veh (hypoxia-exposed, vehicle-treated), 7.5 (hypoxia-exposed, 7.5mg/kg/day clemastine-treated) and Norm (normoxia-exposed) mice. N=3 mice/condition. NS, not significant, ANOVA followed by Tukey’s *post hoc* test.

### Ultrastructural analysis of the minimum effective dose of clemastine

We assessed myelin by electron microscopy (EM) in the corpus callosum of P14 mice in hypoxia-exposed mice treated with vehicle or clemastine at the MED (7.5 mg/kg/day), and in normoxia-exposed mice (Fig. 5a-c). We measured myelin thickness by performing g-ratio analysis, where the diameter of the axon is divided by the diameter of the axon plus the myelin sheath. Clemastine treatment at the MED significantly improved (decreased) the average g-ratio of hypoxia-exposed mice as compared to vehicle treatment, providing ultrastructural evidence that clemastine increases myelin thickness (p<0.0001, ANOVA followed by Tukey’s post hoc test). The average g-ratio in hypoxic mice treated with 7.5 mg/kg/day clemastine was not significantly different from normoxia. These results demonstrate that the MED of clemastine restores myelin thickness in the corpus callosum of hypoxia-exposed neonatal mice to normoxia levels at P14.

**Figure 5.**
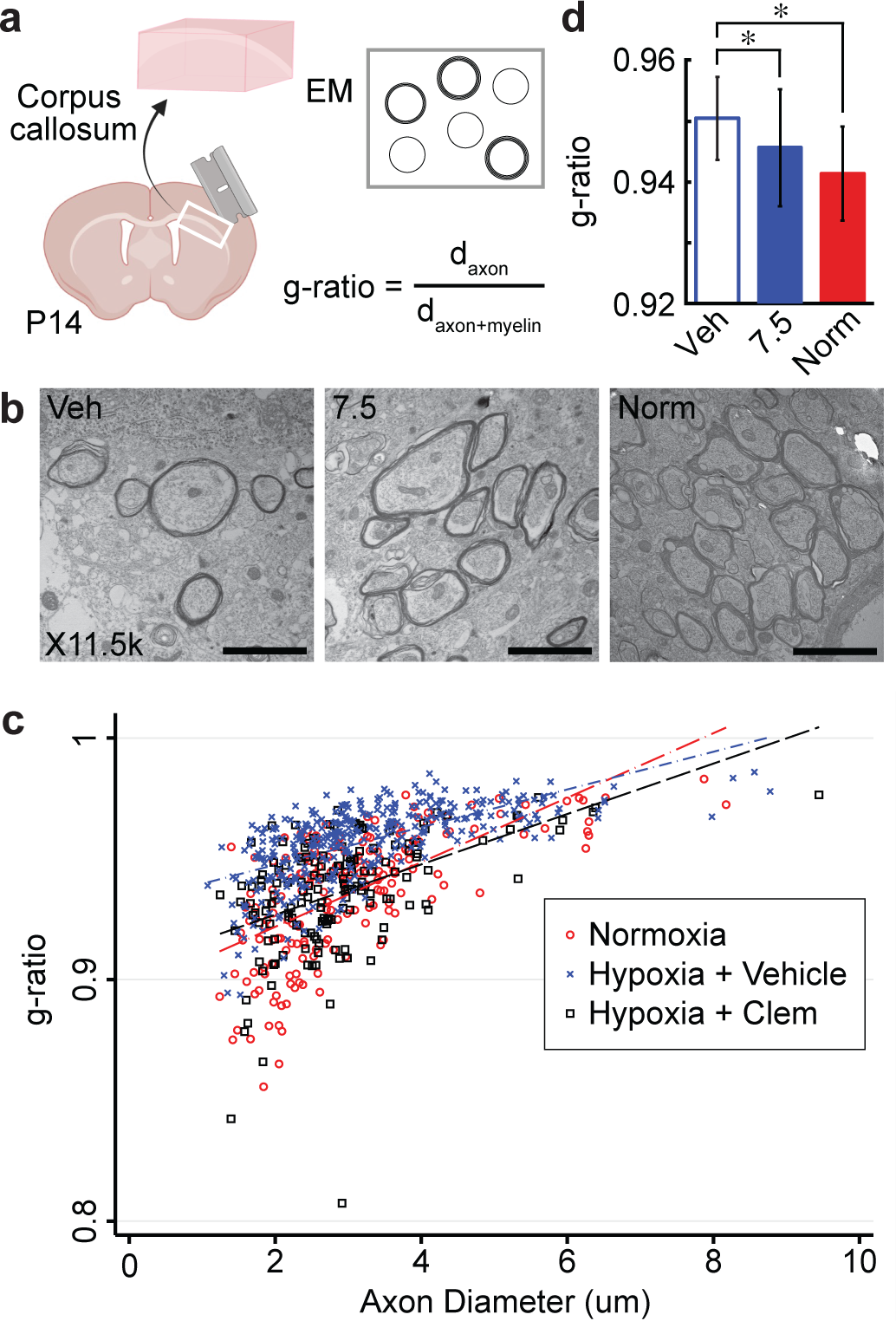
Clemastine restores myelin thickness after chronic hypoxia exposure in neonatal mice. (a) Diagram of experimental setup for g-ratio analysis in P14 mice. EM, electron microscopy; d_axon_, diameter of axon; d_axon+myelin_, diameter of axon plus myelin sheath. Created with BioRender.com. (b) Representative electron microscopy images of the corpus callosum at 11,500x magnification. Scale bars, 2 um. (c) Scatter plot of g-ratios and axon diameters from the corpus callosum at P14, with superimposed linear regressions from normoxia-exposed (red), hypoxia-exposed/vehicle-treated (blue), and hypoxia-exposed/7.5 mg/kg/day clemastine-treated (black, “Hypoxia + Clem”) mice. (d) Comparison between conditions of average g-ratios. Error bars represent S.E.M. Between 530-870 axons were quantified per condition from a total of N=4-8 mice/condition. Hypoxia-exposed animals were: Veh (vehicle-treated) and 7.5 (clemastine 7.5 mg/kg/day). Norm, normoxia-exposed animals. *p<0.05, ANOVA followed by Tukey’s *post hoc* test.

### Pharmacokinetics of the minimum effective dose of clemastine

Neonatal drug development presents many challenges, as neonates often have altered absorption, distribution, metabolism, excretion, and toxicity profiles as compared to adults and older children^33,34^. Similar to human neonates, liver enzyme expression and kidney function undergoes significant postnatal maturation in mice. We sought to determine the pharmacokinetics of oral clemastine in neonatal mice. Fifty mice were treated daily with oral clemastine at the MED, 7.5 mg/kg, starting at P3. Mice were treated for one week to allow plasma concentrations to reach steady-state levels. Treated mice did not appear sedated and did not otherwise have any overt behavioral abnormalities. Mice were sacrificed at planned intervals before and up to 24h following the 8^th^ dose of clemastine for measurement of plasma clemastine concentrations by LC-MS/MS (Fig. 6). The plasma PK parameters are listed in Table 1. The area under the plasma concentration curve (AUC_24_) was 280.1 ng*hr / ml. The elimination half-life (t_1/2_) was 4.6 hours, which is shorter than the t_1/2_ of 21.3 hours previously published in adult humans and more similar to the elimination profile reported in dogs and horses^35–37^. The short half-life resulted in low concentrations (0.7 to 1.4 ng/ml) at the end of the 24-hour dosing interval and suggests that a shorter dosing interval (such as ≤ every 12 hours) could further optimize the pro-myelinating effects of clemastine in rodent models of myelin disorders.

**Figure 6.**
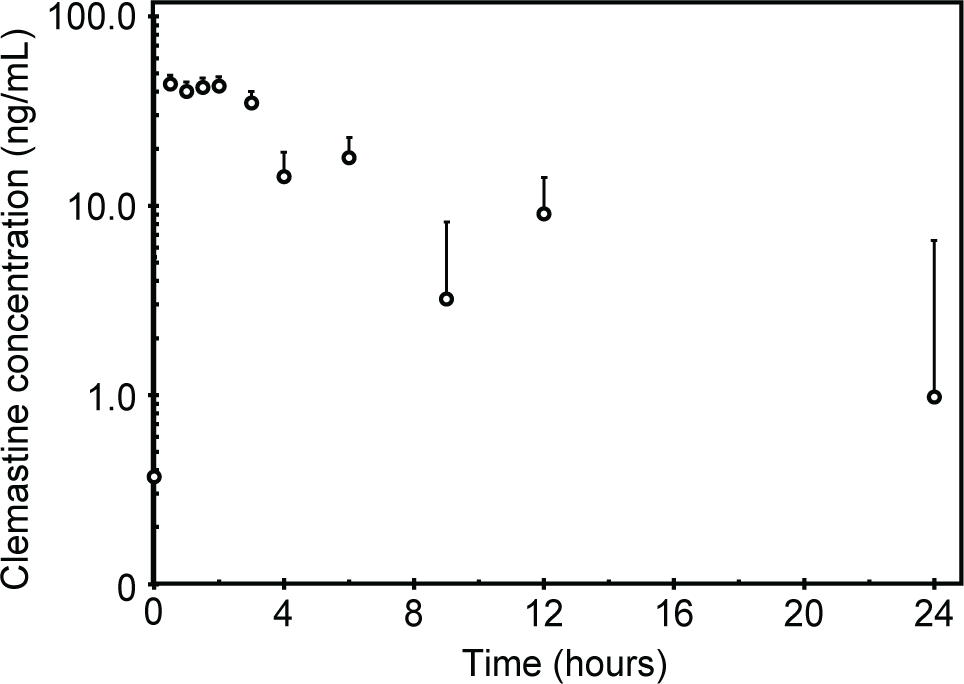
Plasma pharmacokinetics at steady-state of the minimum effective dose of oral clemastine in neonatal mice. Mean (±SEM) plasma clemastine concentration for mice treated with 7.5 mg/kg/day clemastine by oral gavage for one week. N=4-5 mice per time point, with at least one mouse per sex included at each time point.

**Table 1.**
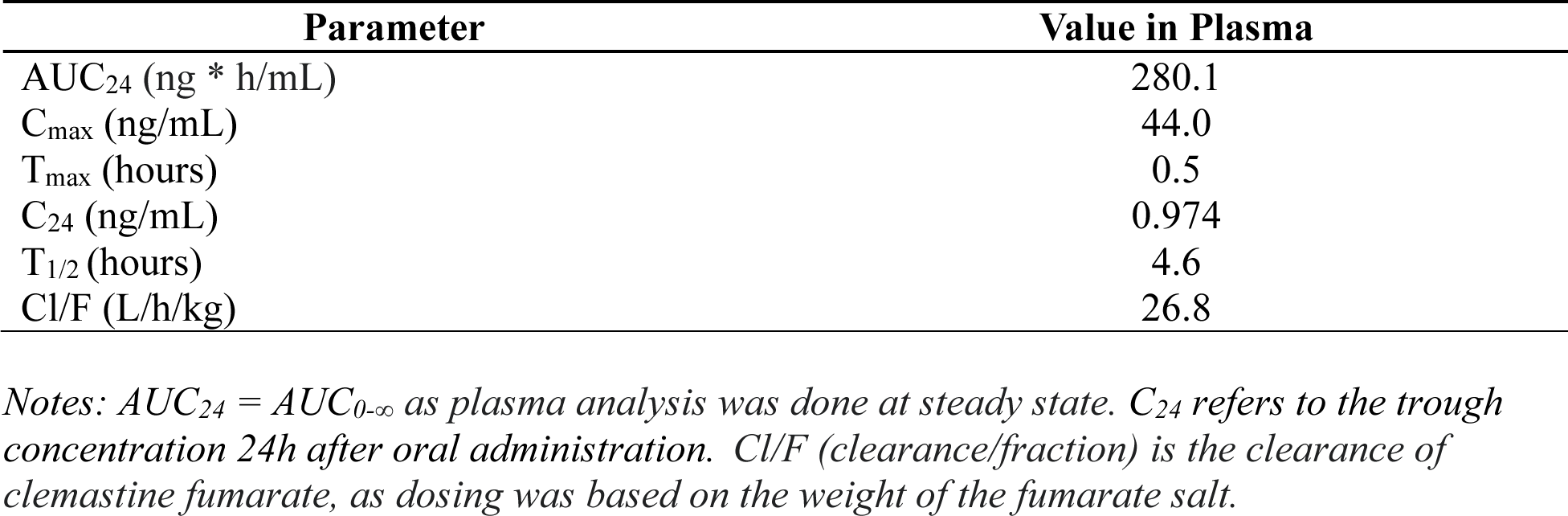
Plasma pharmacokinetics of oral clemastine in neonatal mice.

| Parameter | Value in Plasma |
| --- | --- |
| AUC<sub>24</sub> (ng * h/mL) | 280.1 |
| C<sub>max</sub> (ng/mL) | 44.0 |
| T<sub>max</sub> (hours) | 0.5 |
| C<sub>24</sub> (ng/mL) | 0.974 |
| T<sub>1/2</sub> (hours) | 4.6 |
| Cl/F (L/h/kg) | 26.8 |

*Notes: AUC<sub>24</sub> = AUC<sub>0-∞</sub> as plasma analysis was done at steady state. C<sub>24</sub> refers to the trough concentration 24h after oral administration. Cl/F (clearance/fraction) is the clearance of clemastine fumarate, as dosing was based on the weight of the fumarate salt.*

## DISCUSSION

There are currently no treatments for PWMI that target the oligodendroglial lineage or promote OL differentiation or myelination. Translation of potential treatments from animal models to neonates has been impeded by ethical concerns, drug delivery challenges (such as unavailability or toxicity of liquid oral formulations or lack of blood-brain barrier penetration), and difficulty in predicting PK and toxicity profiles in neonates^38–40^. Furthermore, identification of new therapies for PWMI and other myelin disorders has been slow due to technical barriers to high-throughput screening, such as a requirement for the presence of axons. Here, we demonstrate the MED and associated pharmacokinetics in neonatal mice of clemastine, a myelination-promoting compound identified using a novel high-throughput *in vitro* screening system^12^. Clemastine strongly promotes myelination in multiple settings by directly targeting muscarinic receptors on OPCs^14,15,17–19,29^. Previous mass spectrometry experiments demonstrate that clemastine efficiently penetrates the blood brain barrier and reaches the brain parenchyma^41^ . The medication has demonstrated safety for other indications and is approved in children as young as 12 months old in many countries, with readily available liquid oral formulations. Clemastine may be positioned to overcome some of the challenges that have impeded translation of other potential therapies for PWMI.

Several pharmacokinetic parameters identified in the current study should be considered in future clinical trial design. The CNS-penetrating, pro-myelinating effects of clemastine may require a minimum threshold total daily exposure (AUC_24_), maximum plasma concentration (C_max_), or trough plasma concentration (C_24_). Based on our results, the MED in neonatal mice provides a daily exposure that is approximately three times higher than the previously demonstrated myelination-promoting dose of clemastine in adult humans with multiple sclerosis in the ReBUILD trial^35,42^. Three months of clemastine treatment were required to obtain evidence of remyelination in the ReBUILD trial^42,43^. It is possible that higher daily clemastine doses in patients with multiple sclerosis would allow for a shorter treatment duration or enhanced remyelination. Alternatively, clemastine dosing requirements and ceiling effects may differ in multiple sclerosis as compared to PWMI due to distinct disease mechanisms, patient age, and inhibitory factors such as neuroinflammation. Clemastine has side effects attributed to muscarinic antagonism, such as sedation, that may limit high doses in neonates. Future clinical trials in neonates must optimize daily exposure and plasma concentrations while limiting toxicity and the chance of adverse events.

The current study has several limitations when considering translation to humans. While the murine chronic hypoxia model recapitulates many features of PWMI, hypoxia-induced hypomyelination in mice has been demonstrated to recover over time without intervention^44^. Consequently, chronic hypoxia in mice is thought to cause predominantly a delay in myelination, as opposed to permanent inhibition of OL differentiation. It is therefore possible that hypoxia-induced white matter injury in mice may be more amenable to recovery, and more responsive to myelination-promoting agents, than PWMI in humans. Additionally, we focused our study on assessment of myelination and OL maturation and did not examine behavioral outcomes in clemastine-treated mice. However, improvement in cognitive and motor outcomes after hypoxia has been previously demonstrated after one week of clemastine treatment at 10 mg/kg/day^19^. Given the similarities in histological outcomes between the two doses in our study, we would expect behavioral outcomes to be similar in both dose groups. Finally, while we observed no overt signs of toxicity in mice treated with clemastine at any dose, we did not perform formal toxicity studies or determine the maximal tolerated dose in neonatal mice. Overall, this study is a critical step towards development of a targeted treatment for PWMI. By identifying and characterizing the MED of clemastine that promotes recovery of hypomyelination in a murine chronic hypoxia model, we have overcome a significant barrier to initiation of a well-designed Phase I clinical trial in neonates with PWMI.

## DATA AVAILABILITY

The data generated during the current study are available from the corresponding author on reasonable request.

## ACKNOWLEDGEMENTS

We thank members of the Ostrem, Green and Chan laboratories for technical and conceptual assistance and advice, Drs. Yvonne Wu and Donna Ferriero for insightful discussions and input, and Ivy Hsieh for assistance with electron microscopy experiments.

## AUTHOR CONTRIBUTIONS

B.O. conceived the project. E.O., N.J., and S.R. performed the majority of the experiments with guidance and support from B.O. The pharmacokinetic experiments were designed by A.F. and B.O. and A.F. performed the pharmacokinetic modeling. U.C. and B.S. developed and executed the LC-MS/MS assay and sample analysis. J.C. and A.G. contributed to overall experimental design and data interpretation. E.O. and B.O. wrote the manuscript. All authors critically revised the manuscript and approved the final version.

---

**Supplementary Figure 1.**
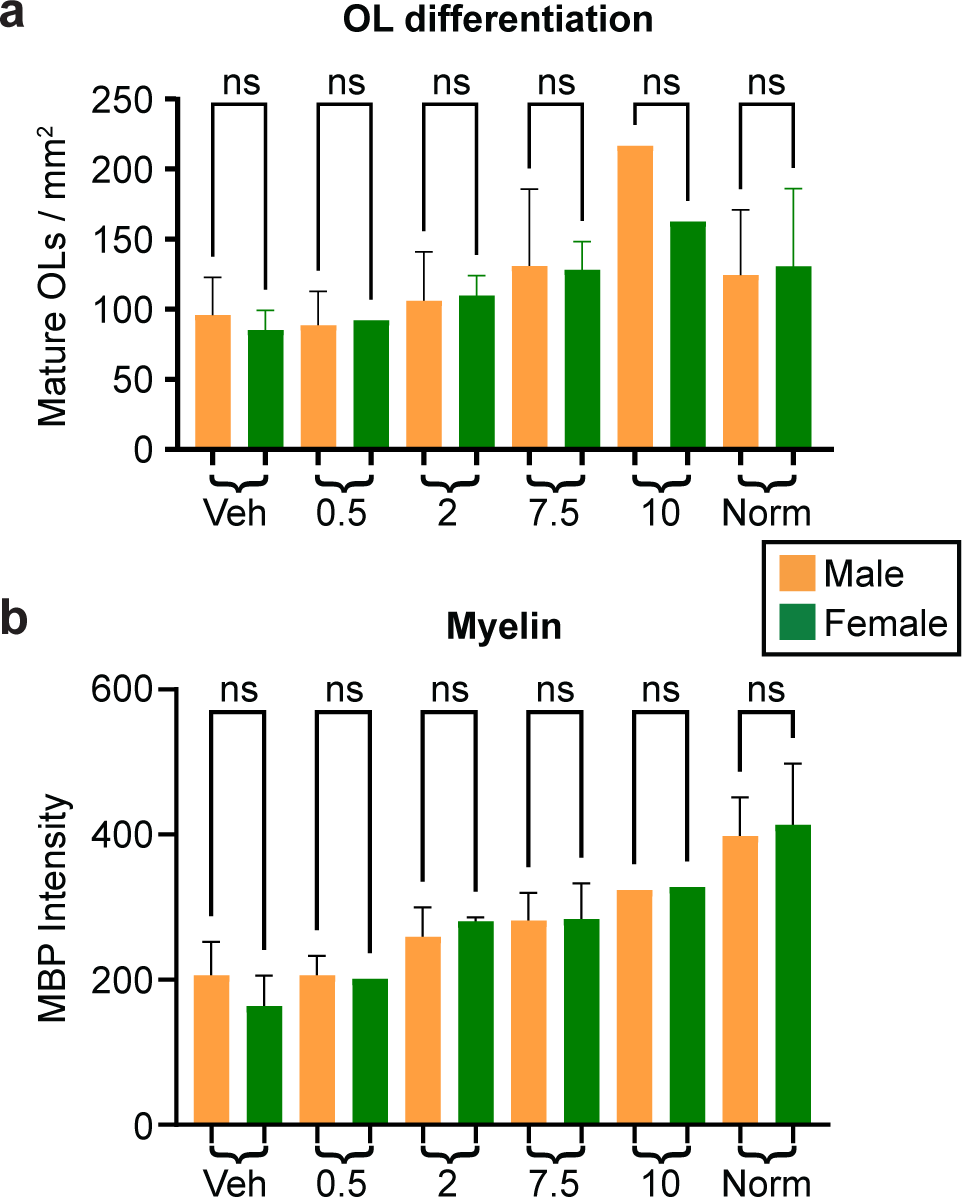
No sex differences in the response to clemastine in hypoxic neonatal mice. (a) Oligodendrocyte density after hypoxia and clemastine treatment, split by sex. N= at least 2 mice/sex/condition. (b) Myelin basic protein (MBP) intensity in the striatum in response to clemastine treatment, split by sex. N= at least 2 mice/sex/condition. No significant differences were observed between males and females of any group, using ANOVA followed by Tukey’s *post hoc* test. NS, not significant. For both graphs, error bars represent standard deviation. M, male (orange bars). F, female (green bars)

